# On the automaticity of visual statistical learning

**DOI:** 10.1101/2022.07.04.498716

**Authors:** Kevin D. Himberger, Amy S. Finn, Christopher J. Honey

## Abstract

Humans can extract regularities from their environment, enabling them to recognize and predict sequences of events. The process of regularity extraction is called ‘statistical learning’ and is generally thought to occur rapidly and automatically; that is, regularities are extracted from repeated stimulus presentations, without intent or awareness, as long as the stimuli are attended. We hypothesized that visual statistical learning is not entirely automatic, even when stimuli are attended, and that the learning depends on the extent to which viewers process the relationships between stimuli. To test this, we measured statistical learning performance across seven conditions in which participants (N=774) viewed image sequences. As task instructions across conditions increasingly required participants to attend to relationships between stimuli, their learning performance increased from chance to robust levels. We conclude that the learning observed in visual statistical learning paradigms is, for the most part, not automatic and requires more than passively attending to stimuli.

## Introduction

The possibility of learning the structure of our environment via mere exposure is alluring. Of course, people learn and remember more when they actively employ a learning strategy (1) or are given an explicit goal to identify associations (2). But some learning processes are thought to be ‘automatic’, in the sense that they can occur incidentally, unconsciously, and without interfering with other processing (3–9). Indeed, the study of statistical learning began with the striking observation that infants can segment repeating sequences of random syllables after only a few minutes of exposure (10), and both children and adults exhibit such learning even if sounds are played in the background while participants are doodling (11). Thus, as originally conceived, statistical learning is a process which ‘*proceeds automatically as a byproduct of mere exposure*’ (12).

Though the consensus is that statistical learning is automatic (4,8,9), the conditions necessary for automaticity have changed over time, especially as statistical regularities have been presented in new modalities and using new cover tasks. In contrast with early statistical learning studies (Saffran et al., 1999), later auditory and visual statistical learning studies indicated that learning could only proceed automatically once participants selectively attended to the stimuli (Turk-Browne et al., 2005; Toro et al., 2005). The literature therefore suggests that, once attention is deployed, visual statistical learning should occur incidentally and without interfering with other cognitive processes (3). In sum, statistical learning, conceived as an automatic process, should proceed even when participants are pursuing a task that is orthogonal to the statistical regularities.

Experimental study of visual statistical learning (VSL) has been strongly influenced by the ostensible automaticity of the learning process. If visual statistical learning is automatic, then a diverse set of cover tasks can be used to measure it, as long as they all require attention to the stimuli. Indeed, the methods for presenting visual regularities have ranged from passive exposure (13,4,14) to 1-back tasks (6,15–17), to tasks involving motion detection (18), image categorization (7,19), background change detection (20), image detection (21), or even direct instruction about the regularities (22).

Motivated by recent demonstrations of confounds in experimental paradigms for measuring automatic learning (23,24), we hypothesized that the learning measured in visual statistical learning experiments is not automatic. Specifically, we proposed that the learning in visual statistical learning paradigms would be heavily influenced by participants’ overt task goals (beyond attention to the stimuli), and the extent to which these goals encourage processing of the relationships between stimuli. Across six statistical learning conditions and one control condition, measuring learning both indirectly and directly, we observed that visual statistical learning did not occur automatically. Instead, participants’ overt task goals strongly modulated learning. Thus, it appears that selective attention to the stimuli is necessary, but not sufficient for robust learning. Instead, the learning in visual statistical learning paradigms is strongly modulated by the extent to which participants are overtly processing the relationships between stimuli.

## Materials & Methods

### Participants

Participants (n=774, ages 18-74) performed visual sequence learning in one of six statistical learning conditions (‘Attend Jiggle’, ‘1-back’, ‘Attend Stimuli’, ‘Attend Relations’, ‘2-back’, ‘Attend Triplets’) or a reference learning condition (‘Feedback Training’). Demographic information is provided in S1 Fig. We analyzed data from 120 participants in each of the 6 statistical learning conditions, and 54 participants in the ‘Feedback Training’ condition. Data were collected online using Amazon’s Mechanical Turk platform, coordinated using PsiTurk (25). Participant compensation varied between $4.50 and $5.75, depending on the approximate duration of each condition (S1 Table). Participation was limited to ‘US only’ participants who had completed at least 99% of their prior Amazon tasks successfully.

### Power analysis

The rationale for our sampling procedure was pre-registered on the Open Science Framework: https://osf.io/yrq8t/. Based on the means and standard deviations in our pilot data we estimated that *a two-sample t-test* could detect an effect in the 2AFC task (see ‘Two-Alternative Forced Choice (2AFC) Task’) with 99% power using 120 participants. The ‘Feedback Training’ condition (n=54) used a smaller sample size given that it served as a reference-point for ceiling performance.

### Exclusion criteria

Because the cover task was a key independent variable in these studies, and because our claims depend on ensuring task engagement, it was critical to ensure high levels of task performance (26) Therefore, we applied a set of 10 pre-registered exclusion criteria to the seven conditions. We collected data from 1,593 participants (774 analyzed, 687 excluded, 132 held-out to balance conditions), with: 4 excluded for data corruption or incorrectly recorded data, 10 excluded for detecting an insufficient number of cover task elements, 92 excluded for detecting an insufficient number of targets (during a target detection post-test), 73 excluded for an excessive number of ‘focus off’ browser events (in which the participant changed focus from the experimental browser tab >20 times), 21 excluded for reporting they could not see all images during the exposure phase, 29 excluded for reporting they had previously participated in a similar experiment, 328 excluded for excessive keypresses (generally >59 keypresses beyond the maximum number of cover task elements (60) or >30 keypresses beyond the maximum number of 2AFC trials (32)), 61 excluded for an insufficient number of 2AFC choices made during a post-test (i.e. fewer than 30 of 32), 68 excluded for reporting they did not understand part of the instructions, and 1 excluded for exceeding the permissible experimental duration (S1 Table). In addition, participants were prevented from participating if their Amazon Mechanical Turk ID had participated in a previous condition or pilot of this experiment.

Under these strict performance criteria, conditional exclusion rates ranged from 29% to 65% (S1 Table). Especially high exclusion rates were seen in the ‘2-back’ condition (261 participants excluded), likely because this is a cognitively demanding task and because the data was collected in the early months of the COVID-19 pandemic. However, the 2-back data are consistent with other conditions collected at other time points and our conclusions do not hinge on the data from this condition.

### Procedure

All conditions consisted of an exposure phase (in which participants were exposed to visual sequential regularities) followed by a test phase (in which we indirectly and directly assessed their knowledge of the regularities).

### Exposure Phase

During exposure, participants viewed a stream of images. The stream was composed of pseudo-randomly ordered ‘triplets’ (3 images in a fixed order). Each image stream employed 4 triplets, composed of 12 distinct images (Fig 1a; Schapiro et al., 2012). The same 12 images were used to generate the image streams across all participants, but the assignment of images to triplets (i.e. the ‘stimulus set’) differed for each participant in each condition. The same collection of 120 ‘stimulus sets’ was used across all statistical learning conditions. Each stimulus set was counterbalanced to ensure that images appeared equally often in each triplet position (i.e. equally often in the first, second, and third position). In the statistical learning conditions, image streams were composed of 720 images (i.e. 60 presentations of each triplet); in the Feedback Training condition, image streams were between 24-168 images (depending upon how quickly the participant learned; see ‘Feedback Training’ below). Images in all but one condition (‘2-back’) were shown for 800ms with a 200ms inter-stimulus interval (ISI). In the ‘2-back’ condition, we lengthened the ISI to 1200ms, as our pilot data suggested participants struggled with the 2-back task at the faster rate (S1 Table).

**Fig 1:**
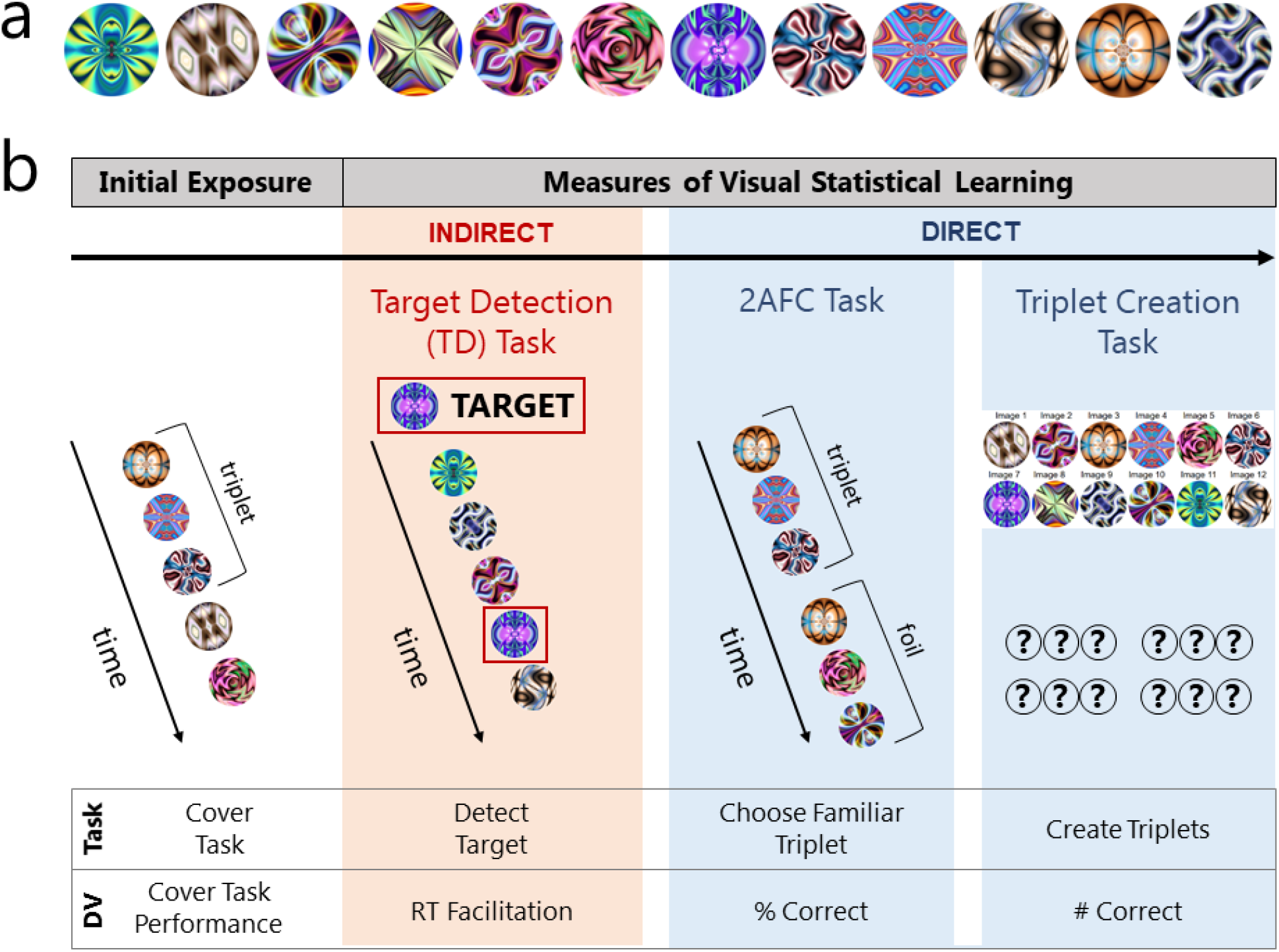
Experimental Paradigm. (a) Experimental stimuli (from Schapiro et al., 2012). (b) Generic structure of conditions. All conditions used a cover task to expose participants to statistical regularities, then assessed regularity learning using three separate measures (1 indirect and 2 direct). ‘DV’ indicates the dependent variable for each phase of the experiment.

In each condition, we manipulated participants’ attentional and goal states during image sequence exposure by assigning them to perform a cover task. Assignments to cover tasks was a between-subjects manipulation, and each participant performed exactly one cover task.

#### Conditions

Below is a brief description of each cover task condition and any significant differences between them (Fig 1b, first column). More detailed instructions and experimental code for each task can be found on OSF (https://osf.io/yrq8t/). In brief, the cover tasks were designed to manipulate the degree to which attention was allocated to the relationships between the stimuli.

1. *Attend Jiggle*. Participants were instructed to watch a stream of images and to detect any image movement from side-to-side (i.e., ‘jiggling’; Turk-Browne et al., 2009). Participants were instructed to press the space bar within 600ms of a jiggle occurring. Participants who missed more than half the jiggles within the 600ms window were excluded from the analysis (n=10; S1 Table). Jiggles occurred for 60 of the 720 images in the stream (i.e. every 12 images on average, ranging 6-31 images). Before beginning the task, participants practiced detecting jiggles in two consecutive stages. In the first stage, participants saw the word ‘PRACTICE’ displayed 20 times, using the same timing as the main exposure phase (i.e. 800ms on, 200ms off). On 4 of the 20 instances, the word jiggled and participants were provided feedback on whether or not they responded to the jiggle quickly enough. The second stage was identical, but no feedback was provided. This second stage was repeated until all four practice jiggles were caught in succession.
2. *1-back*. Participants were instructed to watch a stream of images and to detect any images that were repeated back-to-back (Turk-Browne et al., 2005). Participants were instructed to press the space bar within 800ms of a repeat occurring. Repeats occurred approximately every 13 images (7-32). There were 720 images and 60 repeat-images for a total of 780 images. Participants practiced the 1-back task with black geometric shapes in two stages: a first stage with feedback on correct detection, then a second stage without feedback. The second stage was repeated until all three 1-back repeats were caught in succession.
3. *Attend Stimuli*. Participants were instructed to watch a stream of images and to pay attention to each image, as they would be asked questions about them later (4).
4. *Attend Relations*. Participants were instructed to watch a stream of images and to pay attention to each image and when it appears in relation to other images.
5. *2-back*. Participants were instructed to watch a stream of images and to detect any images that were repeated after one intervening image (Vickery et al., 2012). Participants were instructed to press the space bar within 2000ms of a 2-back repeat occurring. Repeats occurred approximately every 13 images (8-33), with 60 of the 780 images repeating, as in the ‘1-back’ condition. Participants practiced the 2-back task with black geometric shapes in two stages: a first stage with feedback, then a second stage without feedback. The second stage was repeated until all three 2-back repeats were caught in succession.
6. *Attend Triplets*. Participants were instructed to watch a stream of images where the images occur in groups of three. They were instructed that there would be four such groups, and their goal was to learn the order of the images in each of the four groups (29). Participants performed no practice phase since their task did not require any overt action.
7. *Feedback Training* (i.e. benchmark learning condition with explicit feedback). Participants were instructed that their goal was to learn the image sequences composing four triplets. Participants were told to name the images or come up with other mnemonics to better learn the groupings. Participants were shown each of the four triplets twice, with an intervening 1000ms pause between the first and second stream of triplets. Stimulus timing was matched with the statistical learning conditions. After seeing each triplet twice, participants were asked to re-create all four triplets (see ‘Creation Task’ below). Participants were then provided explicit feedback on their performance. If they successfully created all four triplets, they moved on to the test phase. Otherwise, they were told the number of triplets they had successfully created and they then saw the same exposure phase again (i.e. each triplet shown twice, following by the Creation Task). Participants were limited to 20 presentations of each triplet, but all participants successfully re-created the triplets within 2-14 presentations. The exposure and test phases were otherwise identical to the ‘Attend Jiggle’ condition. Since the ‘Feedback Training’ condition served as a benchmark for explicit learning, performance was expected to be high and fewer participants were tested (54 vs. 120). The first 18 stimulus sets from the statistical learning conditions were used, with each stimulus set seen by 3 participants.

#### Supplemental Conditions

Two additional samples were also collected to provide conceptual replication of the ‘Attend Jiggle’ condition with a different stimulus set (see ‘Minimal learning when the cover task was orthogonal to regularities’). Participants in an online condition were drawn from the Amazon Mechanical Turk participant pool (‘Shapes Online’; n=120, ages 18-74), while an in-lab condition participated for course credit at Johns Hopkins University (‘Shapes Offline’; n=42, ages 18-24). The methodology for these additional samples was identical to the ‘Attend Jiggle’ condition, except for minor changes described in the Supplemental Methods (‘Shapes conditions (In-Lab Validation)’).

### Test phase (measures of learning)

After the exposure phase, all participants completed three tests of learning (Fig 1b): (1) a target detection test; (2) a two-alternative forced choice (2AFC) test; and (3) an explicit re-creation of triplets test. All participants performed the tests in this order, to ensure that the most indirect measure of learning would be attempted before directly probing participants’ knowledge.

#### Target Detection (TD) Task

We measured the speed with which participants detected target images within streams of 7-11 images. These streams contained the same triplet sequences seen during the exposure phase, so that target detection speed might be affected by what participants learned about the sequences during the exposure phase. At the start of each TD trial, a single ‘target’ image (one of the 12 images from the exposure phase) was presented to the participant (Fig 1b, second column). Only one target was shown on each trial and each of the 12 targets was used six times (for 72 trials total). Participants initiated the stream of images by pressing the ‘Enter’ key. Once the ‘Enter’ key was pressed, the target image disappeared. Participants were asked to press the space bar upon detection of the target within the stream of images, which were each presented for 200ms with a 40ms ISI, following previous work (6,30). No target detection practice sessions were performed to avoid any possible reduction in the expression of learning that could result from an increased delay of test or processing of novel stimuli.

The position of each target within each 7-11 image stream was balanced across trials. Following previous work, the target image never appeared in the first or last three positions of a trial (Turk-Browne et al., 2005). Therefore, targets could only appear in stream positions 4-8. To ensure that any stimulus could appear at any stream position, it was necessary that streams did not always begin with an image that was the first item in a triplet. Subsequent images followed the triplet structure.

Each image served as the target twice in stream position 4, and once in the other valid stream positions (5, 6, 7 or 8). By making targets appear more often in stream position 4 than in other positions, we aimed to mitigate any response time speeding arising from having a constant hazard rate (i.e. having an increasing probability with each stream position that the target would appear as the next item) and increased expectancy (31).

Collectively, this procedure ensured that all target images (regardless of whether they were in the first, second or third position of their triplet) appeared equally often at each stream position and that there was no relationship between triplet and stream position. The dependent variable was the response time, which we used to calculate response time facilitation between the first and third triplet positions (see ‘Target Detection Task Analyses’).

#### Awareness Probe

Following the TD task, participants were asked if they ‘noticed any patterns in the stream of images’. If they responded positively, they were asked to ‘briefly describe’ the patterns. Regardless of response, participants then proceeded to the more direct measures of learning.

### Direct measures of learning (Forced Choice and Triplet Creation)

#### Two-Alternative Forced Choice (2AFC) Task

The 2AFC phase began by informing participants that: (1) the images they had seen during the exposure phase had followed a pattern and (2) they would need to make judgements about whether groups of images followed that pattern. Participants were told they would choose between two sequences of three images and to choose the group that seemed more familiar.

Participants began by practicing the 2AFC task. They were told to use the left or right arrow key to indicate which of two 3-image sequences, on the left or right, was ‘more familiar’. On each practice trial, instead of the images seen previously, sequences of three circles were presented on the left, then the right, one image at a time. The timing was matched with the exposure phase. The circles were either left blank (incorrect sequence) or contained text indicating the correct choice for that practice trial (e.g. ‘Choose’, ‘the’, ‘left’). Participants could not proceed past this practice phase until they correctly responded to both practice trials.

On each of 32 2AFC trials, participants chose which of two 3-image sequences ‘felt more familiar’ (e.g. Brady & Oliva, 2008). One sequence was a valid triplet, shown during exposure. The other triplet was a positional foil, created by combining first, second, and third position images from three different triplets seen by that participant (Fig 1b, third column). Participants initiated each trial by pressing the ‘Enter’ key. The first triplet was displayed on the left side of the screen, then there was an inter-triplet delay of 1000ms, and then the second triplet was displayed on the right side of the screen. Triplet type (‘real’ or ‘foil’) was counterbalanced such that each type occurred first and second an equal number of times (i.e. 16 trials displayed real before foil triplets and 16 trials displayed foil before real triplets). Each real triplet was presented an equal number of times as each foil triplet, and each foil-real triplet pair was presented an equal number of times. The dependent variable in this forced choice test was the percentage of correct trials.

#### Triplet Creation Task

In the third, and final, measure of learning, participants were informed that the 12 images seen during the initial exposure had been composed of 4 triplets, with the images within each triplet always following one another in the same order. They were then asked to ‘create’ the triplets, by selecting images they felt constituted a triplet. The 12 unique images were shown in a randomized 2 row x 6 column display with an assigned number from 1-12 (Fig 1b, fourth column). Beneath the image display, 12 drop-down boxes were spatially arranged into four groups of three, also numbered 1-12. Each group of three dropdown boxes served as a placeholder for items in a triplet. Thus, by selecting which image should occur in each dropdown box, participants created and endorsed which images they believed constituted each triplet. Triplet re-creation required both image identity and ordering to be correct. Each image could only be used once across all drop-down boxes, to ensure that all 12 images were assigned to a unique position. All 12 drop-down boxes required an entry before the participant could submit their selections and proceed to the post-experiment questionnaire (see ‘Post-Experiment Questionnaire’ in the Supplemental Methods). The dependent variable was the number of valid triplets re-created.

### Analysis

#### Pre-registration

Prior to running our first condition, we pre-registered hypotheses about the learning we would observe in each of the three measures in the ‘Attend Jiggle’ condition (https://osf.io/yrq8t/). After data collection for the ‘Attend Jiggle’ condition, we pre-registered additional predictions for conditions: ‘Attend Stimuli’, ‘Attend Relationships’, and ‘Attend Triplets’ (https://osf.io/v56zx). Conditions ‘1-back’, ‘2-back’, and ‘Feedback Training’ were not formally pre-registered.

### Descriptive analyses

#### Target Detection Task Analyses

The main dependent variable was ‘response time facilitation’ (RTF), specifically the response time difference between images in the first position of a triplet (i.e. less predictable images) and those in the third position of a triplet (i.e. perfectly predictable images). The observation of a facilitation between triplet positions, especially between the first and third position (RTF1-3), is a standard indirect measure of learning and has shown that later images within a triplet are detected more rapidly than earlier images in a triplet (Turk-Browne et al., 2005; Kim et al., 2009; Musz et al., 2015). This metric was calculated both within and across participants.

#### Response time differences between triplet positions

Comparisons of response times between triplet positions employed the Kruskal-Wallis H test as implemented in SciPy 1.3.2 (32). Since multiple tests were run (one for each condition), we corrected for multiple comparisons by applying a Bonferroni correction as implemented in statsmodels 0.10.2 (33). 95% confidence intervals were estimated using a bootstrap procedure (see ‘Bootstrap Procedure’ in Supplemental Methods).

#### Response time facilitation differences between conditions

Comparisons of response time facilitation between conditions employed the Kruskal-Wallis H test as implemented in SciPy 1.3.2 (Jones et al., 2001). Trend analysis on the facilitation data across conditions was performed using the Mann-Kendall test as implemented in PyMannKendall 1.4.1 (34). We report Mann-Kendall tau for the pre-registered conditions (i.e. ‘Attend Jiggle’, ‘Attend Stimuli’, ‘Attend Relations’, ‘Attend Triplets’, ‘Feedback Training’). 95% confidence intervals were estimated using a bootstrap procedure which generated multiple ‘surrogate’ condition-specific datasets by sampling the observed data with replacement, then used the critical values at 2.5% & 97.5% of the resulting, sorted iterations as the interval (for details see ‘Bootstrap Procedure’ in the Supplemental Methods).

#### Response Time Effect Size Analyses

For Kruskal-Wallis H tests, we converted the H statistic (as a proxy for *χ*^2^) into an approximation of *η*^2^. The *η*^2^approximation is shown in Eq. 1 (35). The result of Eq. 1 multiplied by 100 ‘indicates the percentage of variance in the dependent variable explained by the independent variable’ (Tomczak & Tomczak, 2014). Since this is an estimate, and unlike η^2^, the calculation can sometimes produce a negative number, therefore, we only report values greater than zero.

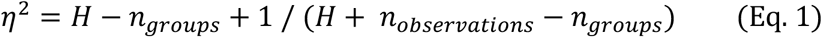

#### Direct Measures of Learning (2AFC & Creation)

The distributions of 2AFC and Creation performance under the null hypothesis were estimated using Monte Carlo simulations (see ‘Monte Carlo Simulation’ in the Supplemental Methods). Since our pre-registered hypotheses anticipated learning, we employed one-way tests to determine critical thresholds in the null distribution. P-values were computed by comparing observed means with critical thresholds in the null distribution. If the overall condition mean exceeded any of the thresholds (p< 0.05, 0.01 or 0.001) we reported the most unlikely threshold exceeded.

##### 2AFC Analyses

Performance on the 2AFC task was assessed by calculating the mean percentage of correct choices made between previously shown, ordered groups of images (i.e. triplets) and group of images that had never been shown in that particular order (i.e. foils).

##### Creation Analyses

Performance in the Creation Task was assessed by calculating the mean number of triplets created. A triplet was considered successfully re-created if all three image identities and their ordering were correct.

#### Bootstrap Confidence Intervals

95% confidence intervals were generated for empirical statistics using a subject-wise bootstrap procedure. We ran 15,000 ‘surrogate experiments’ to generate a distribution of surrogate values for each dependent variable. In each surrogate experiment, we sampled randomly (with replacement) from the experimental data of each condition (e.g. from the list of 2AFC percent-correct scores associated with each participant in the ‘1-back’ condition) to generate a surrogate pool of values, and computed a surrogate mean for that condition. Finally, we sorted the surrogate means from all 15,000 iterations in ascending order, employing the 2.5 and 97.5 percentiles as the boundaries of the 95% confidence interval.

#### Relationships between measures

Spearman rank-order correlations were calculated using SciPy 1.3.2 (Jones et al., 2001).

#### Multiple comparison correction

A Bonferroni correction was applied to all condition-specific p-values. We used a Bonferroni correction factor of 9 to reflect the six statistical learning, one control, and two replication conditions.

## Results

Overall, participants tended to express learning in our direct measures (2AFC & Creation tasks), but not in the indirect measure (i.e. TD task). Therefore, we first focus on the direct measures before discussing the indirect measure.

### Participants recognized and recreated visual regularities

We observed evidence of learning in 6 of 7 conditions using the 2AFC task (Figure 2a) and observed learning across all 7 conditions using the Triplet Creation task (Figure 2b). These learning effects remained statistically significant after correcting for multiple comparisons (all significant p-values <0.001; Table 1).

**Table 1:** Descriptive statistics. All p-values are Bonferroni-corrected. The ‘2AFC % correct by triplets created’ means are only calculated only with sufficient data (4+ participants).

|  |  | **Attend Jiggle** | **1-back** | **Attend Stimuli** | **Attend Relations** | **2-back** | **Attend Triplets** | **Feedback Training** |
| --- | --- | --- | --- | --- | --- | --- | --- | --- |
| **n** |  | 120 | 120 | 120 | 120 | 120 | 120 | 54 |
| **Mean 2AFC (95% CI)** |  | 50.2<br>(48.1, 52.2) | 54.0<br>(51.7, 56.4),<br>p<0.001 | 57.1<br>(54.1, 60.1),<br>p<0.001 | 62.3 (58.6, 66.1),<br>p<0.001 | 66.9<br>(63.3, 70.5),<br>p<0.001 | 66.9 (63.2, 70.7),<br>p<0.001 | 81.3 (76.2, 86.4),<br>p<0.001 |
| **Mean triplets created (95% CI)** |  | 0.13<br>(0.067, 0.20),<br>p<0.001 | 0.15<br>(0.046, 0.25),<br>p<0.001 | 0.32<br>(0.19, 0.45),<br>p<0.001 | 0.74 (0.50, 0.97),<br>p<0.001 | 0.94<br>(0.69, 1.20),<br>p<0.001 | 0.97 (0.72, 1.21),<br>p<0.001 | 3.30 (2.97, 3.63),<br>p<0.001 |
| **Spearman correlation (r<sub>s</sub>; 2AFC vs Creation)** |  | 0.19 | 0.26,<br>p<0.05 | 0.44,<br>p<0.001 | 0.54,<br>p<0.001 | 0.74,<br>p<0.001 | 0.77,<br>p<0.001 | 0.02 |
| **# (%) participants creating x triplets** | **0** | 105<br>(87.5) | 108<br>(90.0) | 94 (78.3) | 84 (70.0) | 69 (57.5) | 66 (55.0) | 2 (3.7) |
|  | **4** | 0 (0) | 2 (1.7) | 2 (1.7) | 14 (11.7) | 17 (14.2) | 15 (12.5) | 38 (70.4) |
| **2AFC % correct by x triplets created** | **0** | 49.1 | 52.6 | 52.9 | 54.9 | 54.8 | 52.8 | n/a |
|  | **4** | n/a | n/a | n/a | 94.1 | 94.5 | 92.1 | 80.6 |

**Fig 2:**
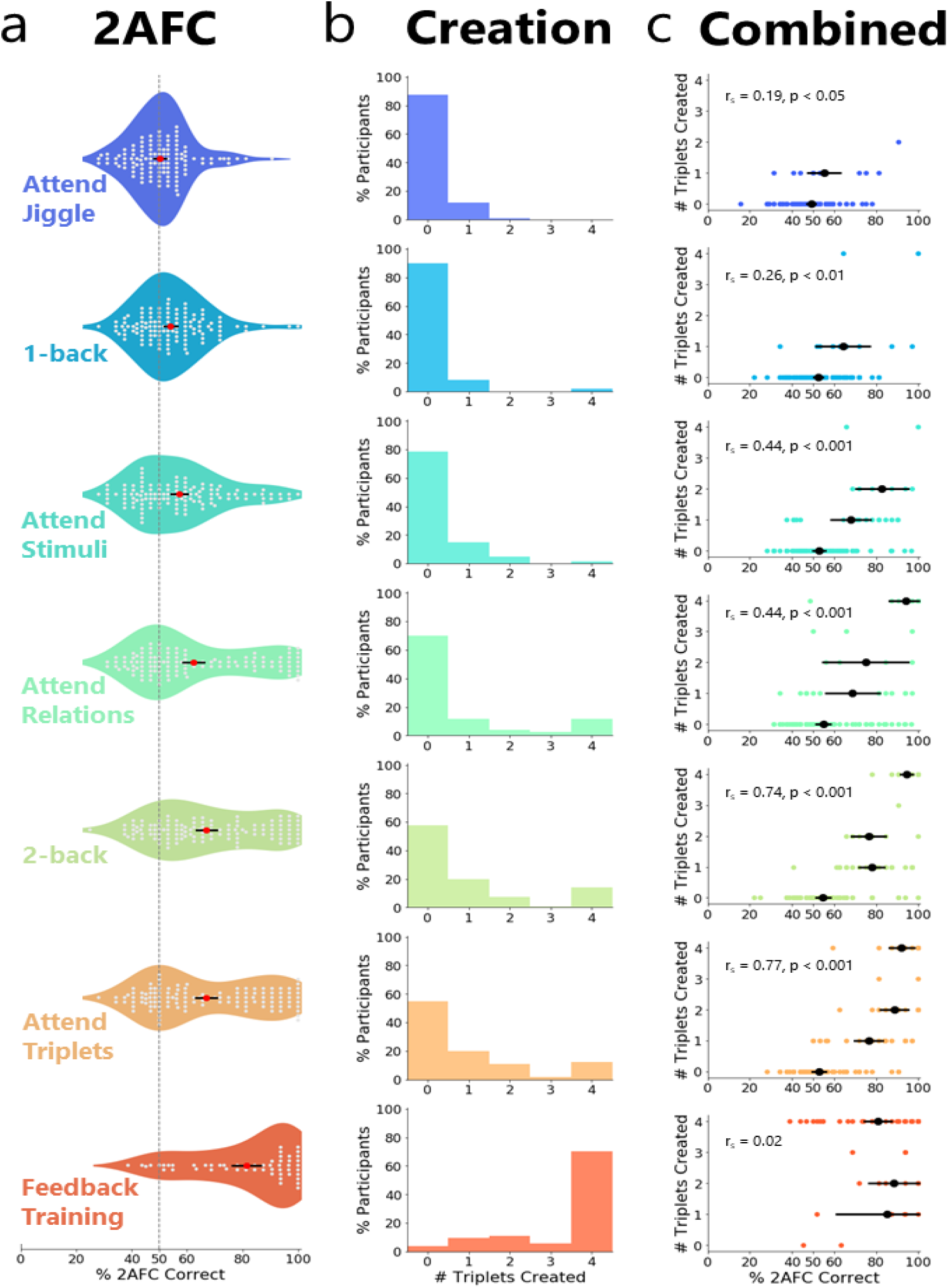
Results from direct measures of learning. (a) Performance of each subject on the 2AFC measure (% of correct choices made), broken down by condition. Each trial has two options, (1) a previously shown triplet or (2) a foil triplet that was never seen. The dashed line indicates chance performance. Each gray dot indicates a participant. The red dot indicates the group mean. (b) Histograms of the number of triplets successfully re-created, by condition. Means and confidence intervals for each condition are reported in Table 1. (c) Relationship between performance on the 2AFC task and the Triplet Creation task. r_s_ indicates Spearman’s rank-order correlation. Each colored dot indicates a single participant. Solid black lines in a & c indicate 95% confidence intervals (where there are at least 4 data points) and black dots indicate group means.

### Attentional and goal states robustly affected learning

Forced-choice familiarity performance was significantly different across learning conditions (2AFC: Kruskal-Wallis H=112.4, p<0.001, η^2^=0.12; Fig 2a). Participants’ ability to re-create the triplets following exposure was also significantly different across the learning conditions (Creation: Kruskal-Wallis H=218.6, p<0.001, η^2^=0.22; Fig 2b).

Given the significant differences between conditions, we ran post-hoc Mann-Kendall trend analyses to test whether cover tasks involving more attention to the relationships (see Methods for ‘Response time facilitation differences between conditions’) were associated with increased learning. In support of our hypothesis, we observed the same increasing trend across conditions in both the 2AFC and Creation tasks (τ=1.0, s=10.0, p=0.027 for both), indicating that increasing attentional demands towards relationships boosted performance in both direct measures of learning.

### Participants who re-created all the triplets performed well on the 2AFC task, while participants who created zero triplets performed near chance levels on the 2AFC task

We observed a significant and positive correlation between the two direct measures (Forced Choice accuracy and Triplet Creation performance) in all conditions except ‘Feedback Training’ (Fig 2c). Additionally, we found that when participants could create all 4 triplets, their average 2AFC performance was near ceiling (Table 1,’2AFC % correct by x triplets created – 4**’**; Fig 2c). This might be expected, as participants who have knowledge of the triplets should be able to exploit that knowledge in the 2AFC task as well. However, statistical learning has long been described as an ‘implicit’ learning process (36). Therefore, we calculated the 2AFC performance amongst those participants who were unable to explicitly re-create any triplets. We found that, for participants who created 0 triplets correctly, their average 2AFC performance was at or near chance (Table 1, ‘2AFC % correct by x triplets created – 0**’**; Fig 2c). Collectively, these data are compatible with the idea that the performance in the Forced Choice task and the Triplet Creation task arise from the same underlying knowledge.

### Noticing a pattern in the exposure stream was correlated with improved forced-choice performance, which increased in tasks requiring attention to relationships

When the task instructions required more attention to the sequential relationships between stimuli, more participants explicitly noticed patterns in the exposure stream (S1 Table). Moreover, those participants who did not endorse ‘noticing a pattern’, showed reduced learning in the 2AFC and Creation tasks relative to those participants who did endorse noticing a pattern (S1 Table). Across all our task conditions, the number of participants who noticed a pattern during exposure was correlated with 2AFC and Creation task performance (S3 Fig).

Noticing a pattern is not equivalent to verbalizable knowledge of the triplet structure. Out of the 600 participants in the five statistical learning conditions where participants were naïve to the four triplets (i.e., excluding the ‘Attend Triplets’ and ‘Explicit Feedback’ conditions), 127 participants endorsed the statement that they had noticed a pattern (S1 Table). Though about 50% of the noticers reported ‘repeated patterns’, ‘order’, ‘sequences’, ‘pairs’, and/or ‘threes’ (n=61; written responses can be found on OSF), only 3 participants were able to verbalize that there were four triplets.

### Minimal learning when the cover task was orthogonal to regularities

While robust evidence of learning was observed in the majority of statistical learning conditions, it was not observed in the 2AFC measure within the ‘Attend Jiggle’ condition (μ=50.19, CI=[48.1, 52.2]). This was surprising, as the dominant view holds that regularity learning occurs as long as selective attention is deployed to the stimuli composing the triplets.

To test the generalization of this finding, we examined two variants of the ‘Attend Jiggle’ condition: one employed a different stimulus set (‘Shapes Online’) and another used the different stimulus set and tested in-lab participants (‘Shapes Offline’). The new stimulus set (S2a Fig) was chosen for comparability with earlier studies (e.g. Fiser & Aslin, 2002). These additional conditions enabled us to test both the generalization of these results across stimulus sets (‘Fractals’ vs ‘Shapes’) and across populations of participants (‘Online’ vs. ‘Offline’).

We found weak evidence of learning in the ‘Shapes Online’ condition (‘Shapes Online’: μ=52.2, CI=[49.8, 54.6], p=0.028], consistent with our observation using the fractal stimuli (Table 1). We observed a 3.1% increase in forced-choice accuracy when testing the shapes stimuli with an in-lab sample (‘Shapes Offline’: μ=55.3, CI=[49.8, 60.9], p<0.001), consistent with prior observations that in-lab samples are comparable to online samples when using proper task controls (37). Quantitatively, participants in the ‘Shapes Offline’ condition demonstrated better performance in the direct measures relative to ‘Shapes Online’ condition (S1 Table); both ‘Shapes’ conditions were better learned than the ‘Fractals’ stimuli in the direct measures (Table 1; S1 Table). Across the three conditions that employed the ‘Jiggle’ cover task (‘Attend Jiggle’, ‘Shapes Online’, ‘Shapes Offline’), there was no significant difference in 2AFC performance (H=2.70), though there was in Creation performance (H=16.9, p<0.01). Further, there was no statistical difference in cover task performance (S1 Table), suggesting that participants attended to the stimuli similarly across conditions.

Overall, there was little evidence of learning in conditions using the ‘Jiggle’ cover task (Table 1; S1 Table). The three ‘Jiggle’ conditions accounted for three of the four lowest levels of 2AFC accuracy, and performance in the Creation task was similarly poor (Table 1; S1 Table). The performance level in the ‘Jiggle’ conditions was so low that the learning effects in these conditions could plausibly be explained by very small subsets of the participants who explicitly learned the regularities (see ‘Which kinds of learning do we measure in standard statistical learning paradigms?’ & S3 Fig). Altogether, we observed little evidence of learning for the majority of participants in the ‘Attend Jiggle’ conditions, despite presenting the same 3-item sequences 60 times over the course of 12 minutes.

### Measures of response time provide no evidence of learning

Response time facilitation effects between first and third triplet positions (RTF1-3) were not consistently observed and differed significantly between the seven conditions (Kruskal-Wallis H=16.5, p=0.011, η^2^=0.013; Fig 3; S1 Table; S7 Fig). Across conditions, three showed numerically negative facilitation and four showed numerically positive facilitation (Fig 3; Table 2). We observed no response time facilitation trends between conditions after conducting a post-hoc Mann-Kendall trend analysis (τ=-0.14, s=-3.0, p=0.76).

**Table 2:** Selected statistics for the response time data.

|  | **Attend**<br>**Jiggle** | **1-**<br>**back** | **Attend**<br>**Stimuli** | **Attend**<br>**Relations** | **2-**<br>**back** | **Attend**<br>**Triplets** | **Feedback**<br>**Training** |
| --- | --- | --- | --- | --- | --- | --- | --- |
| **Mean response**<br>**time facilitation**<br>**1-3 (ms) (95%**<br>**CI)** | 3.19<br>(-4.42,<br>10.8) | -5.09<br>(-13.3,<br>3.13) | 1.13<br>(-8.46,<br>10.7) | 9.85 (-0.36,<br>20.1) | 12.4<br>(3.05,<br>21.8) | -4.74<br>(-14.0,<br>4.54) | -10.7 (-23.1,<br>1.82) |
| **Response**<br>**times by**<br>**triplet**<br>**position**<br>**(1,2,3) (ms)** | 412.2,<br>412.8,<br>409.1 | 439.1,<br>441.6,<br>443.7 | 428.9,<br>435.7,<br>428.2 | 428.6,<br>422.3, 419.9 | 439.3,<br>431.0,<br>428.1 | 427.8,<br>421.4,<br>432.7 | 411.5, 422.9,<br>421.7 |
| **Kruskal-**<br>**Wallis**<br>**difference**<br>**between**<br>**triplet**<br>**positions (H)** | 1.37 | 1.61 | 6.48 | 1.97 | 7.17 | 5.05 | 4.28 |

**Figure 3:**
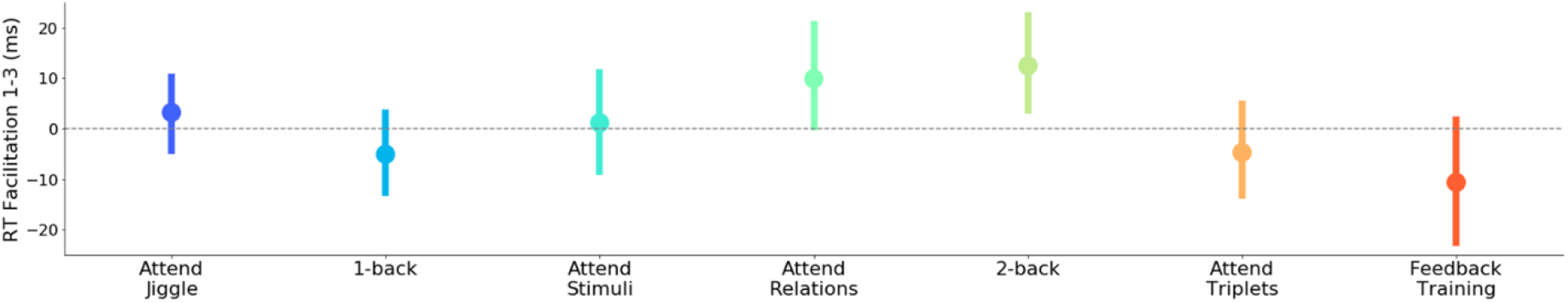
Performance on the indirect measure of learning, response time facilitation between the first and third triplet positions. Line segments indicate 95% confidence intervals.

There was little evidence that more predictable images were detected faster than less predictable images. The two largest RTF1-3 effects were observed in the ‘Attend Relations’ condition (uncorrected p = 0.039, corrected p = 0.35) and the ‘2-back’ condition (uncorrected p = 0.028, corrected p = 0.25). Importantly, our largest response time facilitation of 12.4ms is less than half the size of the previously reported facilitation effects (e.g. 35ms (21), 38ms (38), and 46ms (14)). The RTF1-3 effects in the two conceptual replication conditions were also numerically negative (i.e. in the opposite direction from the effects predicted by the literature) (S2d Fig; S1 Table). Finally, the facilitation effects did not exhibit any correlation with our other measures of learning (S4 Fig), including the ‘Shapes’ conditions.

## Discussion

The visual statistical learning literature suggests that if participants are exposed to sequences containing simple repeated regularities, and if they selectively attend to the stimuli, then visual statistical learning will occur (4,6,39). This is consistent with how ‘automatic’ learning is thought to proceed: operating continuously and unintentionally, and arising unconsciously and without interfering with other processing (3). Our data suggest that visual statistical learning, as measured using conventional methods, is not automatic, because it does not consistently and robustly arise from mere exposure, even when the stimuli are attended. Instead, we found that attentional demands and goal states strongly modulated participants’ learning (Figs. 2a-b), with learning increasing when participants attended to the relationships between stimuli.

The variation in performance across conditions suggests that a major driver of learning was the extent to which participants attended to the relationships between stimuli. We found little evidence of learning in the direct measures (2AFC and Creation) when participants attended to the stimuli but performed a task that was orthogonal to the regularities (e.g. ‘Attend Jiggle’, Figure 2). Conversely, we observed robust learning in the direct measures when the task required participants to compare and relate stimuli across time (e.g. ‘2-back’, Figure 2). Moreover, we found little-to-zero evidence of learning in any condition as assessed by response time facilitation (Figure 3). These data suggest that the learning process, assessed across diverse visual statistical learning paradigms, does not operate equivalently across these paradigms, nor automatically. Rather, despite the commonality that participants attended to the stimuli in all our conditions, almost all the measurable learning could be eliminated (or induced) by changing the task that participants performed with the stimuli.

An alternative explanation to attention to regularities producing the observed learning is that there is a component of learning that is automatic, but augmented by cover task. This would also explain why we observed different levels of learning. This explanation cannot be ruled out, but we emphasize that (i) regardless of condition, the 2AFC performance was at or near chance for participants who could not explicitly re-create any of the triplets (Table 1) and (ii) we observed no learning via the indirect response time task (Table 2). Thus, if there is an automatic component, it does not seem likely that this process was the primary contributor to forced-choice or reaction time effects reported in the visual statistical learning literature.

Visual and auditory statistical learning studies have suggested that selective attention is required to process the relevant stimuli, but that once that occurs, the sequential regularities can be learned without awareness or intent (Turk-Browne et al., 2005; Toro et al., 2005). However, we observed little evidence of statistical learning when participants performed the ‘Jiggle’ cover task, even though participants were attending to the stimuli (Figs. 2a-b, see also Turk-Browne et al., 2009). We therefore propose that cover tasks discouraging relational processing will eliminate or greatly reduce visual statistical learning effects, and that deploying selective attention is not sufficient for learning.

Although it remains unclear how much of participant’s knowledge in statistical learning paradigms is explicit, both during exposure and at test (Ellis, 2009; Arciuli et al., 2014; Batterink, Reber, Neville, et al., 2015; Batterink, Reber, & Paller, 2015; Otsuka et al., 2016), our data suggests that a small subpopulation of participants with explicit knowledge at test could account for most of group-level learning effects that we observed here. For example, in the ‘Attend Jiggle’ condition, there was a single participant (of 120) who was able to successfully recreate all of the triplets. Removing this participant decreases the mean 2AFC performance for this condition below 50%. Moreover, most participants were unable to recreate any triplets in the Creation task, and the 2AFC performance amongst such participants (who could not re-create a triplet) was less than 55% in all conditions (Table 1). Conversely, the best-performing participants on the forced choice test were often those who could also ‘recreate’ the triplets, indicating the knowledge was at least partially explicit (Figure 2c). Thus, a minority of participants with explicit knowledge (possibly arising from idiosyncratic approaches to the task) can account for almost all of the learning in these paradigms (S3 Fig; see also de Leeuw, 2016). More generally, regardless of the cover task, participants who lacked explicit knowledge of the visual regularities demonstrated very little evidence of learning overall (Table 1).

In light of the evidence of explicit processes in statistical learning paradigms, we should examine how the learning processes in statistical learning differ from those that operate in associative memory paradigms (Schlichting et al., 2017) and list-learning paradigms (Howard & Kahana, 2002). Memory in these more conventional learning settings is thought to arise from processes that bind consecutive items to one another and to an unfolding temporal context. Importantly, these associative learning processes can operate incidentally, and the associations require only a single presentation (43,44). The same processes supporting learning in traditional memory paradigms could plausibly explain some memory performance (e.g. forced-choice recognition) in statistical learning experiments, especially memory for sequentially adjacent items. To distinguish statistical learning from other associative learning processes, it will be critical to directly contrast them using common stimuli and participants (e.g. Zhou et al., 2020). Future work should also more precisely manipulate participants attention to and from the stimuli, as well as to and from their sequential relationships.

While we observed lower 2AFC performance than some studies (e.g. Fiser & Aslin, 2002; Schlichting et al., 2017), our results are consistent with others (16,19). Given that the specific items participants learn are affected by properties of the cover task and motor demands during exposure (Vickery et al., 2018) and that attentional demands and goal states impact overall learning (Fig 2), differences across studies could arise from small changes in participants’ task orientation or expectations during the exposure phase.

Why was there no learning evidence in the response time facilitation metric? Our reaction time facilitation paradigm differs from a standard paradigm in previous work (Turk-Browne et al., 2005; Kim et al., 2009; Campbell et al., 2012; Musz et al., 2015; Bays et al., 2015) because our paradigm allows target-detection trials to begin not only with the first element of a triplet, but also with the second or third. Response time speeding effects of items in the second and third triplet positions in prior work (ostensibly a marker of learning) may instead result from a positional confound: that response times decrease for targets that occur later within test trials (23,24).

Our results do not imply that automatic learning is impossible, but rather that the learning we have been measuring in standard visual paradigms is, for the most part, not automatic. There is strong evidence in the auditory domain for automatic mechanisms of regularity detection (e.g. Chait, 2020) and automatic and implicit statistical learning (e.g. Batterink, Reber, Neville, et al., 2015; Aslin, 2017). Humans must (and do) gradually re-organize their internal models in response to visual statistics, and some of these processes must surely be automatic (analogous to perceptual learning), but we question whether we have been measuring such automatic processes in the laboratory.

It may be fruitful to shift to continuous performance paradigms (or ‘online’ tasks, e.g. Siegelman et al., 2018) for measuring statistical learning. In such paradigms, participants perform a task in relation to each stimulus, and their performance for that item (e.g. response time) varies depending on whether the item is consistent or inconsistent with sequential regularities. Continuous performance tasks may reduce the influence of explicit knowledge and task strategies. However, because the ‘encoding’ and ‘testing’ of regularities are interwoven in continuous performance paradigms, the behavioral effects (e.g. response time facilitation) might arise from a single recent exposure to the regularities during the stream. For example participants can develop response time facilitation as soon as the second presentation of an auditory triplet (Batterink, 2017). In such settings, it can be difficult to distinguish learning from short-term priming or retrieval from working memory. Additionally, if continuous performance is to be used as a dependent variable for statistical learning, then we should aim for a standardized metric and task, as response times measured in the context of category-decision, 1-back, and other cover tasks may not be equivalent.

Overall, we find that learning is powerfully modulated by the attentional demands and goal states of the participants in the study. Because directing participants to process the relationships between stimuli shifted their learning performance from chance levels to robust levels, it seems unlikely that we have been measuring an entirely automatic learning process in conventional statistical learning studies. Instead, learning may have been driven by processes that have are also at play in more conventional paradigms such as paired associate learning (45). Altogether, the data prompt us to reconsider how we extract regularities from the world around us, and the range of memory and attentional systems that contribute to this critical learning process.

## Acknowledgments

We thank Chaz Firestone, Mariam Aly, and Anna Schapiro for their insightful feedback and discussion. We also thank Eduardo Sandoval for his assistance on in-lab testing.

## Data Availability Statement

The majority of the conditions were preregistered (Attend Jiggles, Attend Stimuli, Attend Relationships, Attend Triplets) and can be found at https://osf.io/v56zx. De-identified data, analysis scripts, and materials are all available at https://osf.io/vtmpb.

