## Supplementary material for "On the automaticity of visual statistical learning": Supp Appendix

### Supplemental results

#### Fewer foil triplets and more real triplets were created across cover tasks

Since all learning tasks were run in the same order (target detection task, 2AFC, creation), there is a possibility that participants learned the triplet structure during the 2AFC (but not the exposure phase) by chance.  If this occurred, then the number of foil triplets created should be equal to the number of real triplets created (as they are equally likely to be observed by chance). Though more real triplets were created in each condition, we did observe similar numbers of foil and real triplets created in the “Attend Jiggle” and “1-back” conditions, but following the gradient indicated by the post-hoc Mann-Kendall trend analysis, the gap between foils and real triplets created increased with each subsequent condition (S5 Fig; 1)).

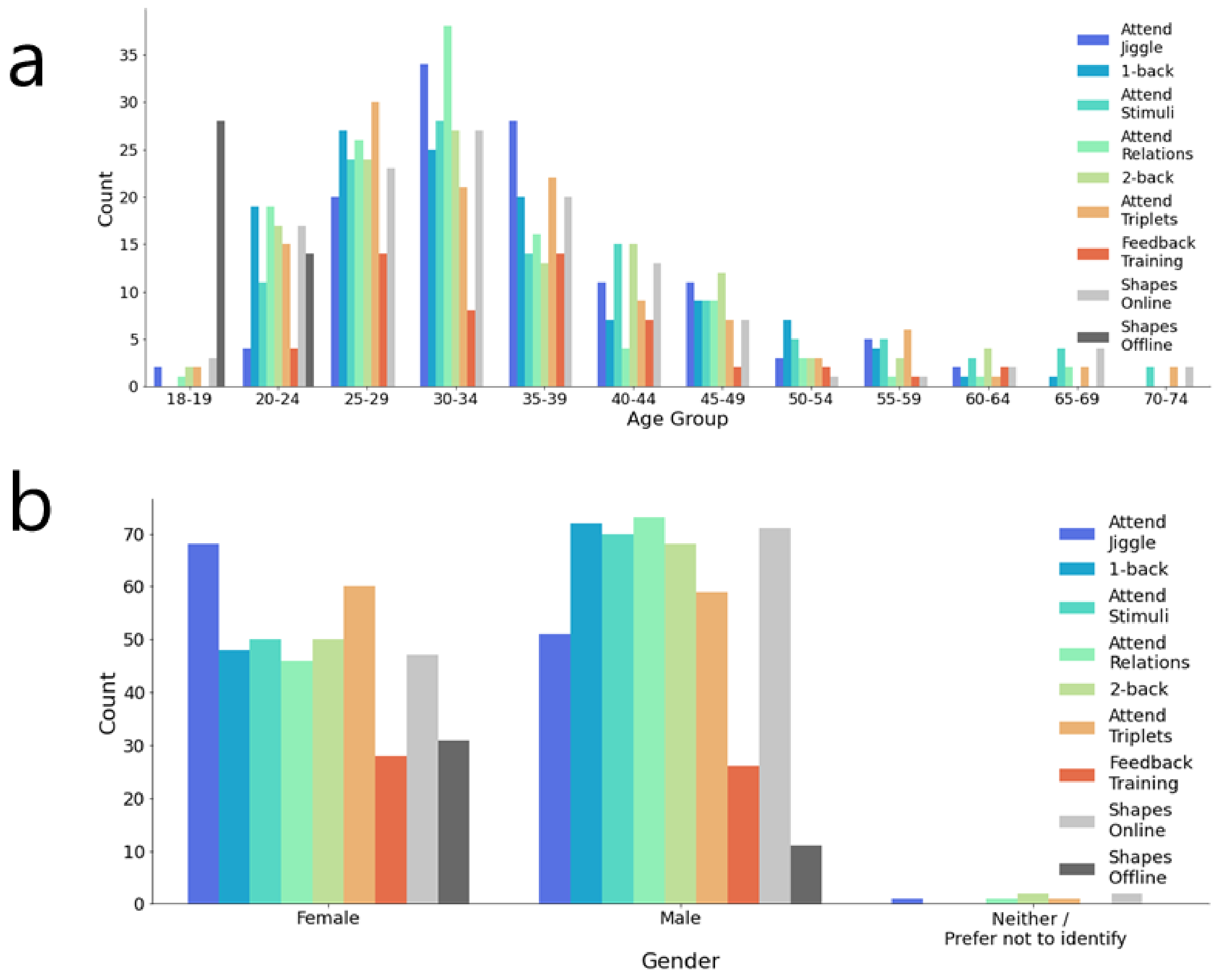


**S1 Fig: Age & gender demographics by condition**. (a) Distribution of age groups by condition. (b) Distribution of gender by condition.

**S1 Table: Experimental parameters, exclusion criteria, and additional statistics on cover task performance and the target detection task**.

| | Attend Jiggle | 1-back | Attend Stimuli | Attend Relations | 2-back | Attend Triplets | Feedback Training | Shapes Online | Shapes Offline |
|---|---|---|---|---|---|---|---|---|---|
| Cover Task | Detect image motion | Detect repeat images | Attend to stimuli | Attend to relationships between stimuli | Detect repeat images | Attend to triplet structure | Explicit triplet training | Detect image motion | Detect image motion |
| # Analyzed | 120 | 120 | 120 | 120 | 120 | 120 | 54 | 120 | 42 |
| Stimulus Type | Fractals | Fractals | Fractals | Fractals | Fractals | Fractals | Fractals | Shapes | Shapes |
| # Excluded -- Total | 51 | 50 | 74 | 96 | 261 | 86 | 69 | 50 | 74 |

| | | | | | | | | | |
|---|---|---|---|---|---|---|---|---|---|
| # Excluded – Data Corrupted or Recorded Incorrectly | 0 | 0 | 1 | 1 | 0 | 1 | 1 | 0 | 1 |
| # Excluded – Insufficient Cover Task Elements Caught | 10 | 0 | n/a | n/a | 0 | n/a | 0 | 0 | 0 |
| # Excluded – Insufficient Targets Caught | 4 | 13 | 10 | 29 | 8 | 19 | 9 | 13 | 10 |
| # Excluded – Excessive "Focus Off" Events | 7 | 7 | 8 | 11 | 14 | 19 | 7 | 7 | 8 |
| # Excluded – Could Not See All Images | 3 | 4 | 1 | 4 | 2 | 1 | 6 | 4 | 1 |
| # Excluded – Previously Participated in Similar Experiment | 3 | 3 | 1 | 3 | 8 | 2 | 9 | 3 | 1 |
| # Excluded – Excessive Keypresses | 4 | 11 | 25 | 35 | 209 | 35 | 9 | 11 | 25 |
| # Excluded – Insufficient 2AFC Choices | 4 | 8 | 21 | 6 | 8 | 3 | 11 | 8 | 21 |
| # Excluded – Did Not Understand Instructions | 16 | 4 | 7 | 6 | 12 | 6 | 17 | 4 | 7 |
| # Excluded – Experimental Duration Exceeded | 0 | 0 | 0 | 1 | 0 | 0 | 0 | 0 | 0 |
| # Not Analyzed (i.e. "Held Out") | 13 | 19 | 28 | 23 | 19 | 18 | 12 | 19 | 28 |
| Compensation | $4.50 | $4.50 | $5.00 | $5.00 | $5.75 | $5.00 | $4.40 | $4.50 | $4.50 |
| Length of Image Stream During Cover Task | 720 | 720 | 720 | 720 | 720 | 720 | Variable (between 24-168) | 720 | 720 |
| Cover Task Stimulus Duration (ms) | 800 | 800 | 800 | 800 | 800 | 800 | 800 | 800 | 800 |
| Cover Task Inter-stimulus Interval (ms) | 200 | 200 | 200 | 200 | 1200 | 200 | 200 | 200 | 200 |

| | | | | | | | | | | |
|---|---|---|---|---|---|---|---|---|---|---|
| 2AFC Stimulus Duration (ms) | | 800 | 800 | 800 | 800 | 800 | 800 | 800 | 800 | 800 |
| 2AFC Inter-stimulus Interval (ms) | | 800 | 800 | 800 | 200 | 1200 | 200 | 200 | 200 | 200 |
| Mean cover task performance | | 90% Jiggles Caught | 83% Repeats Caught | n/a | n/a | 59% Repeats Caught | n/a | 3.07 attempts before all triplets created | 91% Jiggles Caught | 90% Jiggles Caught |
| Mean cover task response time (ms) | | 444 | 526 | n/a | n/a | 742 | n/a | n/a | 447 | 445 |
| Mean percentage of targets caught | | 90% | 86% | 89% | 85% | 89% | 84% | 86% | 92% | 93% |
| Mean target response time (ms) | | 410 | 443 | 430 | 422 | 434 | 426 | 417 | 394 | 381 |
| Mean RTF1-3 (ms) | | 3.2 | -5.1 | 1.1 | 9.8 | 12.4 | -4.7 | -10.7 | -1.6 | -2.9 |
| Mean 2AFC | | 50.2 | 54.0 | 57.1 | 62.3 | 66.9 | 66.9 | 81.3 | 52.2 | 55.3 |
| Mean Triplet Creation | | 0.13 | 0.15 | 0.32 | 0.74 | 0.94 | 0.97 | 3.3 | 0.17 | 0.64 |
| # (%) participants creating x triplets | 0 | 105 (87.5) | 108 (90.0) | 94 (78.3) | 84 (70.0) | 69 (57.5) | 66 (55.0) | 2 (3.7) | 103 (85.8) | 26 (61.9) |
| | 4 | 0 (0) | 2 (1.7) | 2 (1.7) | 14 (11.7) | 17 (14.2) | 15 (12.5) | 38 (70.4) | 0 (0) | 3 (7.1) |
| 2AFC % correct by x triplets created | 0 | 49.1 | 52.6 | 52.9 | 54.9 | 54.8 | 52.8 | n/a | 50.5 | 47.8 |
| | 4 | n/a | n/a | n/a | 94.1 | 94.5 | 92.1 | 80.6 | n/a | 95.7 |
| # Endorsing "Patterns Seen" before 2AFC | | 9 | 19 | 31 | 40 | 28 | 44 | 14 | 13 | 12 |
| RTF1-3 for Pattern Seers (ms) | | 9.25 | -21.9 | -4.09 | 18.0 | 23.3 | 4.03 | -14.1 | -2.93 | -19.5 |
| RTF1-3 for Pattern Unseers (ms) | | 2.70 | -1.93 | 2.95 | 5.78 | 9.13 | -9.82 | -9.45 | -1.41 | 3.73 |
| 2AFC for Pattern Seers | | 53.1 | 54.0 | 66.5 | 78.2 | 79.2 | 76.4 | 84.3 | 50.4 | 62.3 |
| 2AFC for Pattern Unseers | | 50.0 | 54.0 | 53.9 | 54.4 | 63.2 | 61.4 | 80.3 | 52.4 | 52.6 |
| Creation for Pattern Seers | | 0.44 | 0.053 | 0.55 | 1.58 | 1.79 | 1.34 | 3.43 | 0.31 | 0.75 |

| | | | | | | | | | |
|---|---|---|---|---|---|---|---|---|---|
| Creation for Pattern Unseers | 0.11 | 0.17 | 0.26 | 0.33 | 0.68 | 0.75 | 3.25 | 0.15 | 0.60 |
| # Pattern Seers who mention triplet structure | 0 | 0 | 0 | 3 | 0 | n/a | n/a | 0 | 0 |
| # Pattern Seers who mention any structure | 1 | 4 | 16 | 23 | 17 | n/a | n/a | 2 | 5 |
| psiTurk version | 2.1.2 | 3.0.0 | 3.0.0 | 3.0.0 | 3.0.0 | 3.0.0 | 2.2.4 | 2.2.3 | 3.0.0 |

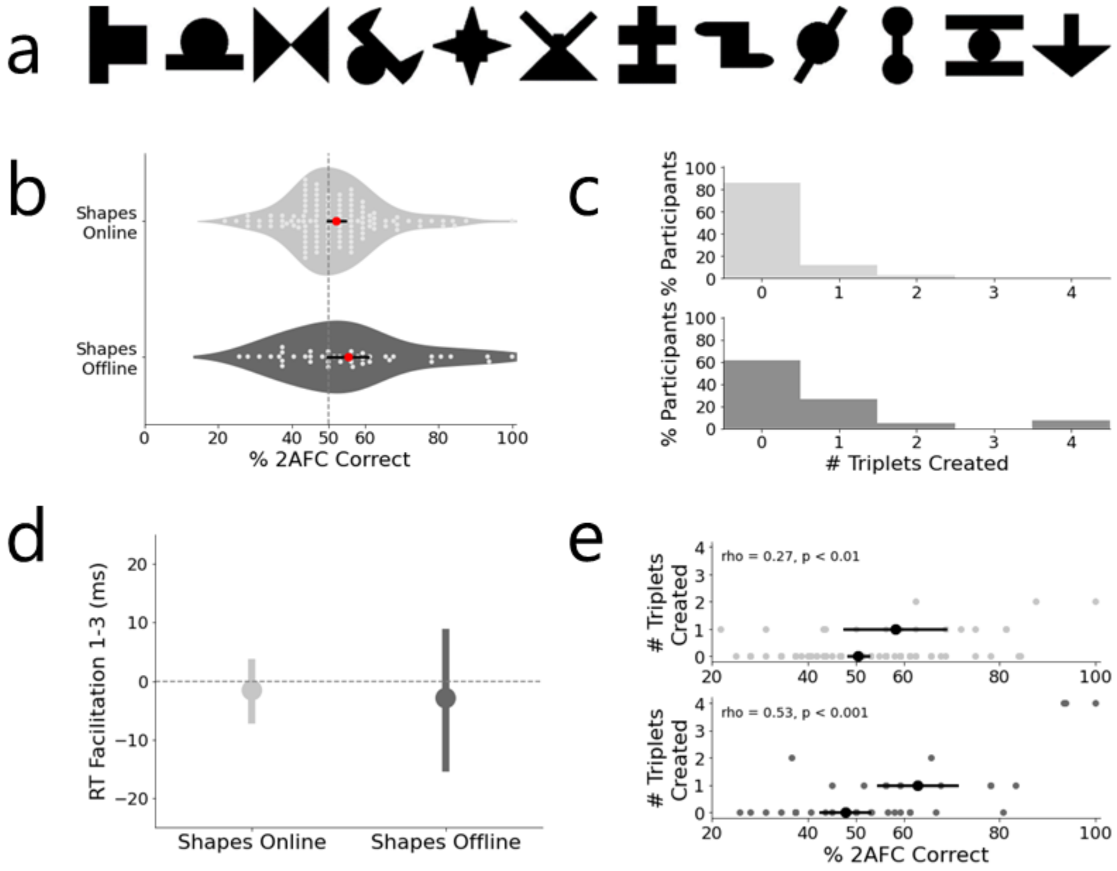


**S2 Fig: Stimuli and results from the "Attend Jiggle" conceptual replications, both in-lab ("Shapes Offline") and out-of-lab ("Shapes Online")**. (a) Stimulus set (b) Distribution and individual performance metrics on the 2AFC task. Red dot indicates mean. (c) Histograms illustrating the percentage of participants creating between zero and all four triplets. (d) Response times in target detection task, split by position of target within a triplet. (e) Response time facilitation between targets in triplet position 1 and triplet position 3. (e) Relationship between performance on the 2AFC task and the Creation task. All line segments convey 95% confidence intervals.

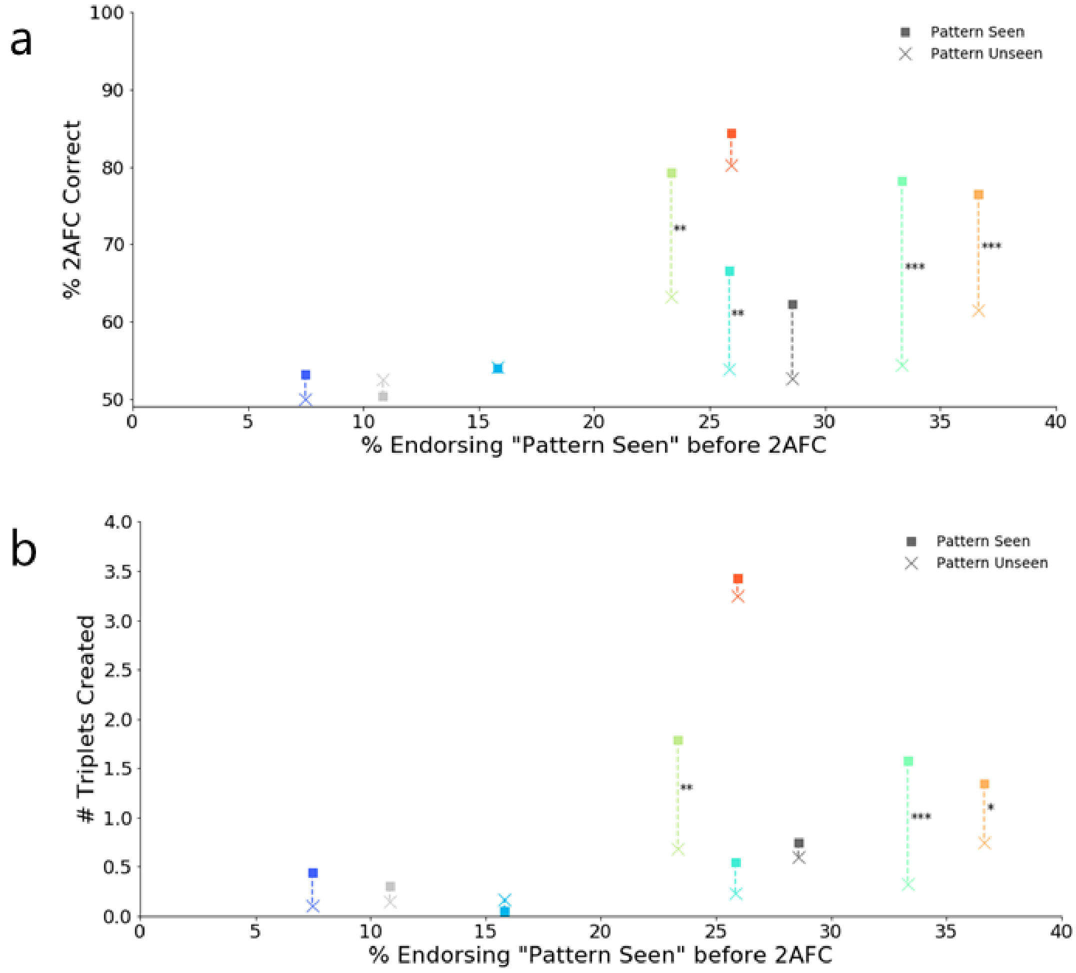


**S3 Fig: Direct task performance by condition when separating participants who "noticed a pattern" and those who did not**. (a) 2AFC task performance. (b) Creation task performance. Asterisks indicate level of significance for difference between means after applying Bonferroni correction: *: p<0.05, **: p<0.01, ***: p<0.001.

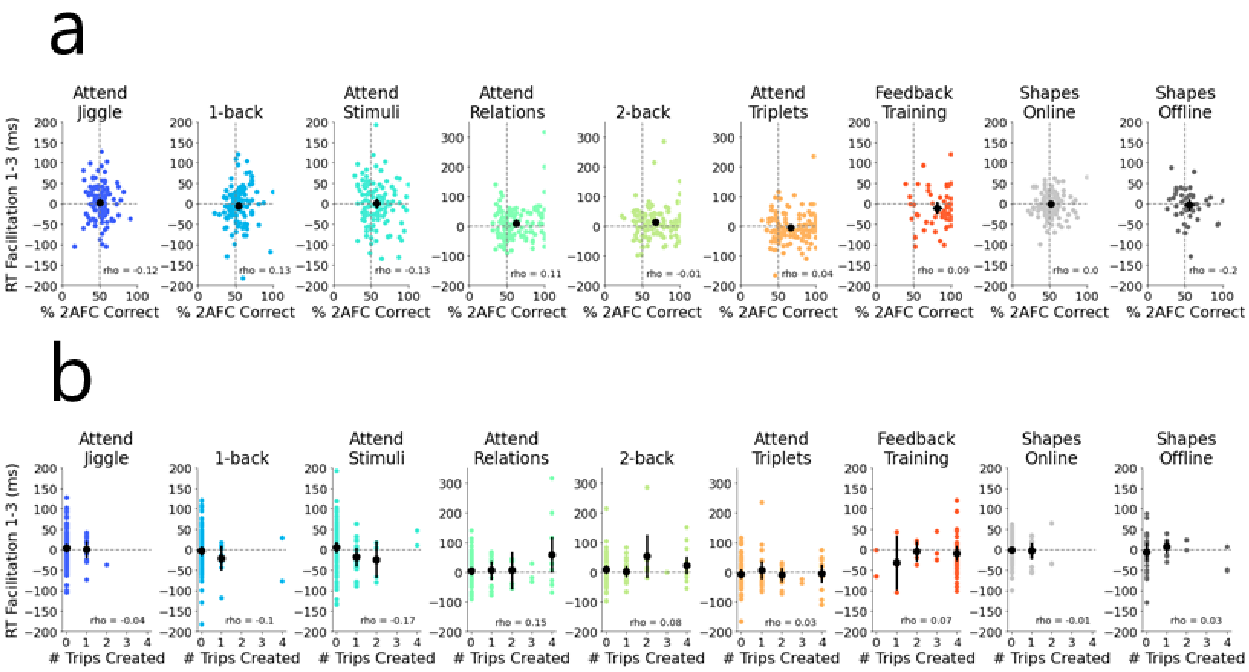


**S4 Fig: Relationships between response time facilitations and the two direct measures**. (a) Relationship between the response time facilitation and percentage of 2AFC trials correctly chosen. (b) Relationship between the response time facilitation and number of triplets successfully created. Black line segments indicate 95% confidence intervals.

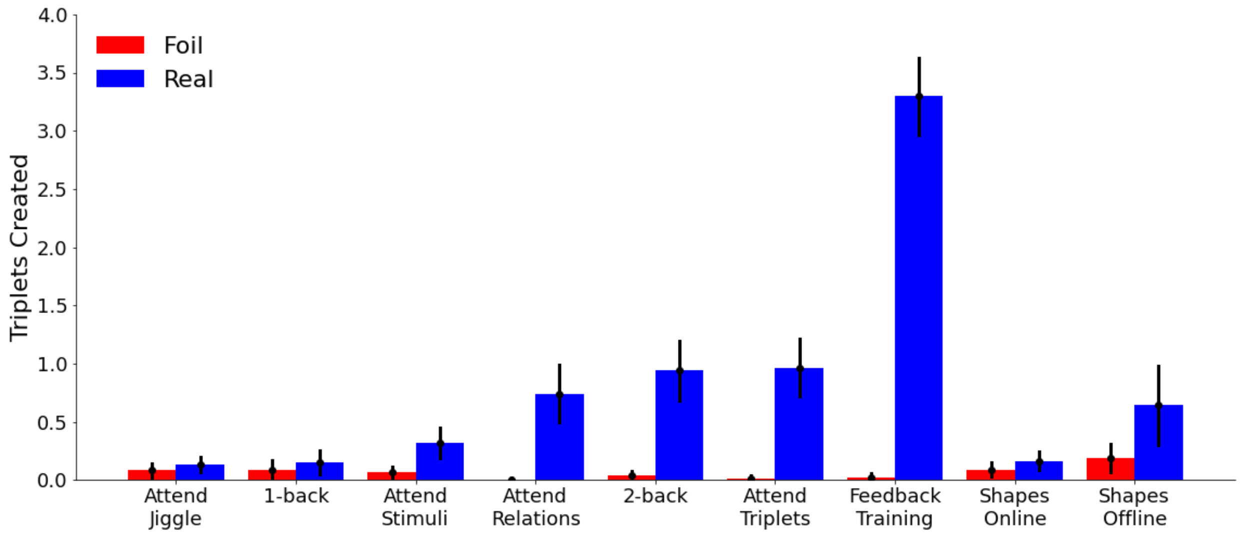


**S5 Fig: Comparison of real and foil triplets created during the creation task**. Each triplet type was seen an equal number of times during the 2AFC task.

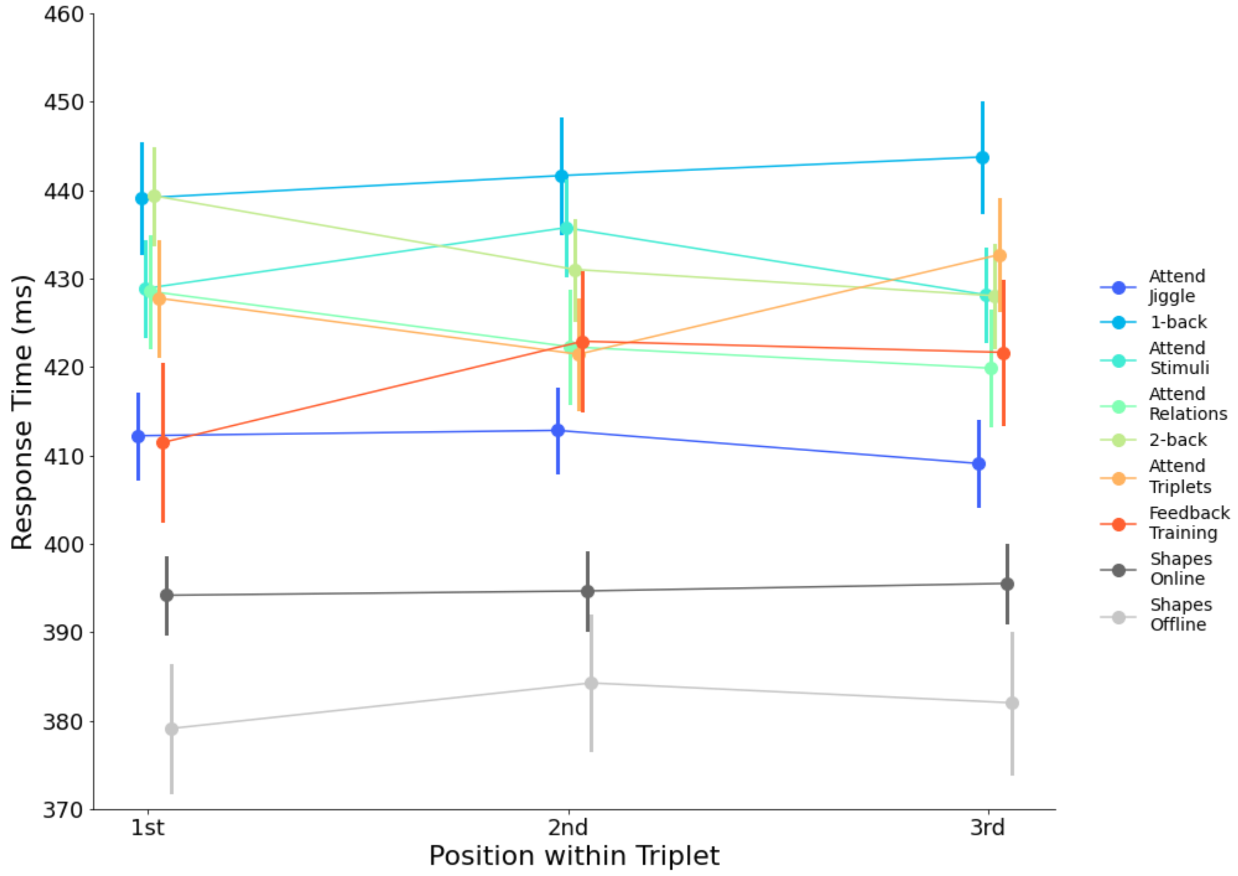


**S6 Fig: Response times across conditions and triplet positions**. Line segments show 95% confidence intervals.

### Supplemental Methods

#### Exclusion criteria added after pre-registration

We implemented new exclusion procedures after the onset of COVID-19 due to decreasing data quality (especially noticed in the "2-back" condition). Data collection preceded the onset of COVID-19 lockdown precautions for "Attend Jiggle", "Attend Stimuli", and "Feedback Training". However, data collection for "Attend Relations" began at the end of February 2020 and “Attend Triplets” data collection began in early March 2020. The “2-back” data collection began in the beginning of May 2020 with “1-back” beginning data collection at the end of May 2020. In brief, we began excluding participants in real-time as opposed to after they completed the full experiment. Participants excluded in this manner were ones that exhibited obviously malicious behavior (e.g. pressing no keys or systematically pressing keys unrelated to the task). This change allowed us to improve data quality in real-time and reduced the cost of running experiments and resulted in a reduced “excluded” count for these conditions.

#### Shapes conditions (In-lab Validation)

We conducted two experiments in the “Attend Jiggle” condition using different stimuli (S2a Fig). We had hypothesized that simpler stimuli (geometric shapes) might improve performance on the direct measures of learning. The first, “Shapes Online” condition was identical to the “Attend Jiggle” condition, except the stimuli used were black geometric shapes (Fiser & Aslin, 2002). The second, “Shapes Offline” condition involved 42 undergraduate students from Johns Hopkins University, participating for course credit. Since the in-lab version had fewer participants, we used only the first 42 stimulus sets from the larger sample (2–5).

#### Post-Experiment Questionnaire

Participants were asked a series of questions about the experiment upon completion of the Triplet Creation task. They provided a self-assessment of their performance on each of the three measures of learning. They were also asked if they knew there were four triplets prior to being told about the triplets in the instructions for the Creation task. They were asked whether they used any strategies in each of the phases. Additionally, they were asked if they had mentally assigned labels to any of the images and, if so, to list any labels they had assigned. The full questionnaire can be found on OSF.

### Simulation & Bootstrapping Analyses

For all simulations or resampling, the parameters were set to equate the simulated/resampled data with the observed data. The number of surrogate participants was equated to the condition being analyzed (n=120, except for "Explicit Feedback" where n=54). The number of simulations and bootstrap iterations performed was 15,000. The number of: stimuli (12), triplets (4), elements per triplet (3), and 2AFC trials (32) were the same for all conditions. Code is available on the Open Science Framework (https://osf.io/vtmpb).

#### Monte Carlo Simulation

Determining whether the observed data was significantly different than chance required estimating a data distribution under the null hypothesis. We used a resampling approach to simulate results for surrogate participants across many surrogate experiments. Once the null distribution was generated, we then determined critical thresholds using our pre-registered hypotheses. We used one-tailed tests for the 2AFC and Creation simulations, given that we expected learning to be above chance. In addition to the critical thresholds for $\alpha=0.05$, we also calculated the critical thresholds for $\alpha=0.01$ & $\alpha=0.001$. If the actual mean exceeded any of three levels of significance ($p<0.05$, $p<0.01$, and $p<0.001$), we reported the most unlikely to have occurred by chance. Otherwise, we reported only the measured value.

***2AFC Task Resampling Procedure.*** An array of length 32 (i.e. the number of forced-choice trials) was populated with half zeros and half ones to represent correct left (0) or right (1) choices. This array was used as "ground truth" for all iterations. For each iteration, a surrogate conditional mean was calculated by first calculating a surrogate subject mean for each fake participant, and then taking the overall mean of all surrogate participants for that iteration. The mean for a surrogate participant was calculated by generating an array of 32 elements (i.e. number of trials) and randomly populating it with zeros and ones. This allowed us to calculate the "number 2AFC correct" for each surrogate participant. By repeating this for 15,000 iterations we derived a null distribution of forced-choice accuracy scores for each condition. We calculated the overall surrogate conditional mean and a critical threshold for "chance" performance, using an alpha of 0.05 (i.e. taking the value at the 95$^{th}$ percentile of the null distribution).

***Creation Task Resampling Procedure.*** This simulation was used to estimate chance levels of performance for the task of triplet creation. We estimated chance of levels of creating genuine triplets and generating foil triplets. An array containing the integers 1-12 (inclusive) were shuffled, with each integer representing one the 12 stimuli presented in each experiment. This randomized array served as the "ground truth" for all iterations of the shuffling procedure for a particular surrogate participant. Virtual triplets were defined by splitting the 12 values into sets of three (i.e. "0", "1", "2" is triplet 1) and then compared against a fixed stimulus set across all surrogate participants. For each surrogate experiment, we calculated the surrogate participant mean by checking how many randomly shuffled triplets match the fixed stimulus set triplets. By repeating this process for 15,000 iterations we generated a null distribution of "creation" scores. From this distribution, we calculated the overall chance creation mean and a critical threshold for “chance” performance, using an alpha of 0.05 (i.e. taking the value at the $95^{th}$ percentile of the null distribution).

#### Counterbalancing procedure for stimulus sets

Counterbalancing was achieved by creating “mini-groups” of three stimulus sets, where the item positions were circularly shifted within a triplet across stimulus sets. Thus, subject X might be assigned the first stimulus set in a mini-group, and would see a triplet composed of the stimuli: ($A_1$, $A_2$, $A_3$). Subject Y might be assigned the second stimulus set in the mini-group, and would see a triplet composed of the stimuli: ($A_2$, $A_3$, $A_1$). Finally, subject Z, would see a triplet composed of: ($A_3$, $A_1$, $A_2$). In this way, each stimulus image, $A_1$ or $A_2$ or $A_3$, appears equally in the first, second and third position of a triplet. This circular shifting procedure ensures triplet position is uncorrelated with the properties of individual stimuli.

#### Statistical packages used

##### Differences in 2AFC performance between “Pattern Seers” and “Pattern Unseers”

Mann-Whitney U tests were conducted using SciPy 1.3.2 (6) with a Bonferroni correction applied using statsmodels 0.10.2 (7) to correct for multiple comparisons.

##### Rank-biserial correleations

The rank-biserial correlation (RBC) and the matched pairs rank-biserial correlation (MPRBC; Kerby, 2014) were calculated for the Mann-Whitney U and Wilcoxon signed-rank tests, respectively, using the Python package Pingouin 2.7 (8).

### Supplemental Discussion

#### How generalizable are these findings to statistical learning generally?

##### Auditory statistical learning

One common difference between visual and auditory statistical learning paradigms is that many auditory learning paradigms rely upon stimuli with which the participants have previous experience (e.g. syllables, tones, chirps). Previous work has shown that stimuli with with which we have more experience are easier to statistically learn (10). In contrast, our stimuli (and visual statistical learning stimulus choices more generally) were chosen to be novel. We hypothesize that using visual stimuli with comparable levels of prior knowledge to syllables might equate the performance on post-exposure measures. In addition to this possibility, it is also possible that auditory statistical regularities are easier to learn in the temporal domain, relative to visual statistical regularities. It is likely that there are modality-specific learning processes for statistical regularities.

##### Spatial paradigms for visual statistical learning

Ngiam et al. (2019) convincingly show that only those participants with explicit knowledge are able to leverage statistical regularities in a spatial array. They argue that this effect is the result of leveraging long-term memory of the statistical regularities. Additionally, they suggest that implicit learning might occur only for spatial paradigms (e.g. Zhao et al., 2013). The work cited in this manuscript did not have spatial components tied to the regularities to-be-learned.

#### Motor paradigms for statistical learning

We focused our literature review on research that did not have motor actions tied to the regularities to-be-learned. Our assumption was that these paradigms involve different neural circuits and are more likely to result in implicit knowledge (13,14).
